# Identification of scavenger receptor BI as a scavenger of free heme that is essential for protection against hemolysis

**DOI:** 10.64898/2026.04.17.718316

**Authors:** Misa Ito, Jianyao Xue, Ling Guo, Dan Hao, Qian Wang, Alexander H. Williams, Chang-Guo Zhan, Ailing Ji, Preetha Shridas, Wen Su, Shu Liu, Zhenheng Guo, Mingcui Gong, Scott M. Gordon, Bin Huang, Jianhang Jia, Chieko Mineo, Philip W. Shaul, Xiang-an Li

**Affiliations:** Department of Pharmacology and Nutritional Sciences, University of Kentucky College of Medicine, Lexington, KY 40536, USA; Saha Cardiovascular Research Center, College of Medicine, University of Kentucky, Lexington, KY, USA; Department of Pharmaceutical Sciences, University of Kentucky College of Pharmacy, Lexington, KY 40536, USA; Department of Internal Medicine, Division of Endocrinology, University of Kentucky College of Medicine, Lexington, KY 40536, USA; Division of Cancer Biostatistics, University of Kentucky College of Medicine, Lexington, KY 40536, USA; Markey Cancer Center, University of Kentucky College of Medicine, Lexington, KY 40536, USA; Center for Pulmonary and Vascular Biology, Department of Pediatrics, University of Texas Southwestern Medical Center, Dallas. TX 75390, USA; Lexington VA Healthcare System, 1101 Veterans Drivel, Lexington, KY 40502, USA; Department of Physiology, University of Kentucky College of Medicine, Lexington, KY 40536, USA

## Abstract

Severe hemolysis is a life-threatening condition with limited therapeutic options. Although haptoglobin and hemopexin sequester hemoglobin and heme, these protective systems are rapidly saturated during acute hemolysis, leading to the accumulation of cytotoxic free heme. In this study, we identify scavenger receptor BI (SR-BI) as a critical mediator of free heme clearance. SR-BI binds heme and facilitates its hepatic uptake under pathological conditions. Mice lacking hepatic SR-BI exhibit impaired heme clearance and increased susceptibility to heme- and hemolysis-induced lethality. Pharmacological upregulation of hepatic SR-BI via imatinib or adenoviral delivery confers protection against heme toxicity. Using a humanized model of sickle cell disease (SCD), we further demonstrate that sickle hepatopathy significantly reduces hepatic SR-BI expression compared to non-SCD littermates, potentially increasing vulnerability to heme-induced injury. Notably, adenoviral-mediated SR-BI upregulation rescues SCD mice from heme toxicity. These findings reveal a previously unrecognized mechanism of heme detoxification via hepatic SR-BI and identify a promising therapeutic target for hemolytic disorders.

**One-Sentence Summary:** Identification of scavenger receptor BI as a targetable scavenger of heme in hemolysis

## Introduction

Severe hemolysis is a life-threatening condition that can occur in malaria, sickle cell disease (SCD), sepsis, extracorporeal circulation and transfusion reactions^1, 2^. Upon hemolysis, broken red blood cells release hemoglobin (Hb), which subsequently releases heme. Hb and heme bind to haptoglobin (Hp) and hemopexin (Hx), forming Hb-Hp and heme-Hx complexes. These complexes are then cleared in the spleen and liver through their receptors, CD163 and CD91 respectively^3–5^. Whereas Hb-Hp-CD163 and heme-Hx-CD91 systems are crucial to scavenge Hb and heme, in cases of severe hemolysis, the Hb-Hp-CD163 and heme-Hx-CD91 systems become saturated due to depletion of Hp and Hx, resulting in high levels of free heme in the circulation. Free heme is highly toxic that causes organ injury and death^6, 7^. While there are several treatment options for managing hemolysis, heme toxicity remains largely an untreated condition. Currently, treatments for hemolysis, depends on etiology, include anti-malaria drugs, hydroxyurea, transfusions, C5 complement inhibitor, plasmapheresis, immunosuppressant, and splenectomy^8^. Despite these efforts aimed at limiting further hemolysis, heme toxicity may still develop in these conditions and result in lethality. Thus, there is an urgent need to address heme toxicity. In an elegant prior study, Tolosano et al showed that unexpectedly Hp/Hx double knockout mice showed better survival in severe hemolysis compared to both wild-type and single-knockout mice, suggesting the existence of other heme scavenger pathways^9^. Identifying novel heme scavengers may reveal new therapeutic targets for heme toxicity during hemolysis.

Scavenger receptor class B type I (SR-BI) is a multifunctional membrane protein that exerts its diverse biological roles through ligand binding^10, 11^. It is abundantly expressed in the liver, adrenal glands, and endothelial cells. In the liver, SR-BI binds high-density lipoprotein (HDL) to facilitate reverse cholesterol transport, a process that contributes to cardiovascular protection^12^. In the adrenal glands, SR-BI mediates the uptake of HDL-derived cholesterol, providing the substrate for glucocorticoid synthesis in response to stress, thereby supporting host defense during sepsis^13^. In endothelial cells, SR-BI-HDL binding enhances nitric oxide production via endothelial nitric oxide synthase (eNOS), promoting vascular homeostasis^14^. Paradoxically, SR-BI also binds and transports low-density lipoprotein (LDL) in endothelial cells, a mechanism that contributes to atherogenesis^15^. Beyond lipoproteins, SR-BI interacts with lipopolysaccharide (LPS), facilitating its uptake and detoxification^16, 17^. Additionally, SR-BI recognizes and binds apoptotic cells, aiding in their clearance^18^.

In this study, we identify hepatic SR-BI as a novel scavenger of free heme. SR-BI binds heme and facilitates its uptake under pathological conditions. Mice lacking hepatic SR-BI exhibit impaired heme clearance and heightened susceptibility to heme- and hemolysis-induced mortality. Conversely, upregulation of hepatic SR-BI via imatinib or adenoviral delivery confers protection against heme-induced lethality. Furthermore, in a humanized SCD mouse model, we observed a significant reduction in hepatic SR-BI expression compared to non-SCD littermates. Importantly, restoring hepatic SR-BI expression via adenoviral delivery rescued SCD mice from heme-induced mortality. These findings reveal a previously unrecognized mechanism of heme clearance mediated by hepatic SR-BI and highlight its potential as a therapeutic target for the treatment of hemolysis-associated heme toxicity.

## Results

### SR-BI binds to heme and facilitates the cellular uptake of free heme

To investigate the binding of heme with SR-BI, we performed computational modeling of human SR-BI based on the homologous LIMP-2 protein^19^. Our model shows that heme binds to the entrance of the hydrophobic cavity of SR-BI receptors with the Gibbs free energy of −111 Kcal/mol, indicating a spontaneous process (**Fig. 1a**). Hemin positions above F195 providing two pi-stacking interactions with the protoporphyrin ring of Hemin. Additionally, a strong hydrogen bond is formed with N330, and a charge-charge interaction is formed between one of the carboxylic acid motifs and a nearby K151. (**Fig. 1b**). The CPC motif (residues 321–323) plays an important role in maintaining this binding environment by enforcing a disulfide-dependent structural constraint that preserves pocket geometry and supports the N330–heme hydrogen-bonding network. Importantly, these sites (residues 151 to 330) are highly conserved between mouse and human (**Fig. S1b**). We also simulated the binding pose for heme fluorescent analog ZnMP (**Fig. 1c**) and the SR-BI inhibitor, BLT-1^20^ (**Fig. S1a**), which show that both ZnMP and BLT-1 occupy the same binding site at the entrance of the SR-BI hydrophobic cavity as hemin.

**Fig. 1.**
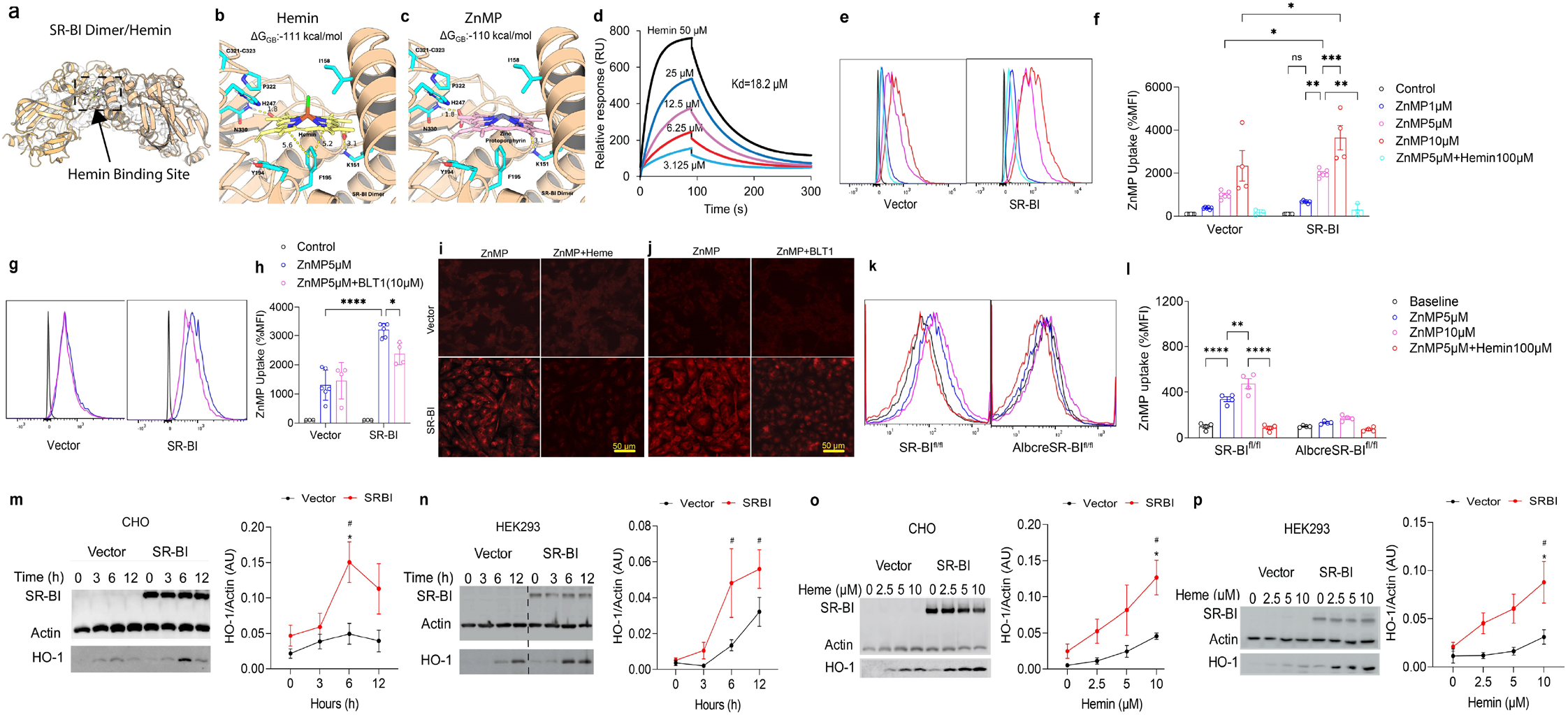
SR-BI binds to heme and facilitates its uptake. (**a**) An overview of the hemin binding site of a SR-BI dimer’s hydrophobic pocket (arrow). (**b**) The binding pose of hemin with SR-BI. (**c**) The binding pose of hemin fluorescent analog ZnMP with SR-BI. (**d**) Dissociation constant was measured via SPR. At 30 μL/min, with association for 90 s, and dissociation for 210 s, the association rate constant was Ka = 1.10 × 10^3^ M^−1^·s^−1^, and the dissociation rate constant was Kd = 2.00 × 10^−2^ s^−1^. (**e-h**) Flow cytometry analysis of ZnMP uptake in CHO cells expressing SR-BI or vector. Control, 1 μM, 5 μM ZnMP (n=6). 10 μM ZnMP (n=4). 5 μM ZnMP + 100 μM hemin (n=3). Representative histograms are shown in (**e**,**g**), and the %MFI representing ZnMP uptake is shown in (**f**,**h**). (**h**) ZnMP uptake in the presence of BLT-1 (10μM). 5 μM ZnMP (n= 6). 5 μM ZnMP + BLT-1 (n=4). (**i-j**) Confocal fluorescence microscopy for ZnMP uptake in the presence/absence of heme and BLT-1. (**k-l**) Flow cytometry analysis of ZnMP uptake in primary hepatocytes from SR-BI^fl/fl^ and AlbCreSR-BI^fl/fl^ mice (n=4). (**m-n**) CHO and HEK293 cells expressing SR-BI or vector were treated with 10 μM hemin for time dependent HO-1 expression. (**o-p**) CHO and HEK293 cells expressing SR-BI or vector were treated with various hemin concentrations for 6hrs. HO-1, SR-BI, and actin expressions were analyzed by immunoblotting. The blots are representative of 3 independent experiments. The * denotes comparison with the control at the same timepoint, while # denotes comparison with baseline. The means ± SEM are plotted. * P < 0.05; ** P < 0.01; *** P < 0.001; **** P < 0.0001; Significances were determined by two-way ANOVA.

Next, we employed surface plasmon resonance (SPR) to measure the binding affinity between human SR-BI and heme. The estimated equilibrium dissociation constant (Kd) was 18.2 μM (**Fig. 1d**). It is worth noting that the average free heme levels in SCD and malaria patients can reach 70 µM^21,22^. Thus, SR-BI can bind heme at pathological conditions.

Then, we tested if heme-SR-BI binding leads to intracellular heme uptake using Chinese hamster ovary (CHO) cells stably expressing SR-BI. We incubated ZnMP, a fluorescent heme analog with uptake kinetics and transporter dependence identical to those of hemin^23–25, 26^, with CHO cells and measured ZnMP uptake by flow cytometry. Intercellular ZnMP was detected in CHO-vector cells, indicating free diffusion of ZnMP into cells as a lipid molecule (**Fig. 1e-f**). CHO-SR-BI showed a significantly higher intracellular presence of ZnMP at 5 and 10 µM, indicating that SR-BI facilitates heme uptake (**Fig. 1e-f**). The uptake of ZnMP was significantly reduced in the presence of a high dose of hemin (**Fig. 1e-f**). SR-BI inhibitor BLT-1 significantly decreased intracellular ZnMP in CHO-SRBI but not in CHO-vector, indicating specific uptake via SR-BI (**Fig. 1g-h**). Next, we aimed to evaluate if SR-BI mutants would have diminished capacity for heme uptake. Specifically, we examined SR-BI C323G. Prior studies, including ours, have shown that this mutant is correctly localized to the cell membrane, yet completely lacks the ability to uptake cholesterol esters from HDL^27, 28^. This loss of function suggests that the C323G substitution disrupts the receptor’s lipid channel and therefore provides a useful model for verifying SR-BI–dependent heme uptake. To investigate how this mutation affects heme binding, we performed in-silico docking simulations on the C323G variant (**Fig. S1a**). Consistent with prior studies, our analyses showed that replacing cysteine with glycine at position 323 eliminates the local disulfide-dependent structural constraint^29^, resulting in destabilization and collapse of the predicted heme-binding pocket. Subsequently, we validated these computational findings using ZnMP uptake assays. CHO cells expressing SR-BI C323G displayed markedly reduced ZnMP uptake, reaching levels comparable to CHO-vector controls (**Fig. S1c**). This data further verified that SR-BI is crucial for heme uptake.

Next, we used confocal microscopy to further confirm heme uptake via SR-BI. Compared to CHO-vector, CHO-SR-BI showed higher intracellular signal of ZnMP, which was reduced in the presence of a high dose of heme and specifically inhibited by BLT-1 (**Fig. 1i-j**). Lastly, we verified the uptake of heme with primary hepatocytes from SR-BI^fl/fl^ and AlbcreSR-BI^fl/fl^ mice. ZnMP uptake was significantly diminished in SR-BI null hepatocytes compared to that of the wildtypes (**Fig. 1k-l**). This result further supports that hepatic SR-BI mediates heme intracellular uptake.

Intracellular heme upregulates heme oxygenase-1 (HO-1) expression, which has been used as a marker for heme uptake^30^. To further characterize the SR-BI-mediated heme uptake, we incubated CHO cells and human embryonic kidney (HEK) 293 cells expressing SR-BI or vector with heme and quantified HO-1 expression at various time points. We observed a greater upregulation of HO-1 expression in CHO-SR-BI and HEK-SR-BI cells than in vector controls, with peak expression of HO-1 at 6 hours (**Fig. 1m-n**). Then, we incubated cells with heme of various doses. We observed greater upregulation of HO-1 expression in CHO-SR-BI and HEK-SR-BI cells than in vector controls, with the highest HO-1 expression at 10 µM of heme (**Fig. 1o-p**). These findings support that SR-BI facilitates the uptake of free heme. To further determine the intracellular fate of SR-BI–imported heme, we co-stained ZnMP with a lysosome marker (Lysotracker). Our images revealed a diffuse cytosolic distribution of ZnMP with minimal lysosomal overlap (**Fig. S1f**), supporting that SR-BI delivers heme to cytosolic/ER-proximal pools relevant for HO-1 activation. Besides being a marker for heme uptake, HO-1 can also be induced due to serum depravation and membrane damage due to lipid peroxidation by heme. While HO-1 expression was induced via these unspecific mechanisms in the vector control cells, cells with SR-BI expression showed significantly higher HO-1 expression at the same timepoint and hemin concentration. Consistent with our confocal imaging results where ZnMP was diffusedly distributed across ER-proximal site that is relevant for HO-1 induction, the robust upregulation of HO-1 supports a specific mechanism rather than a nonspecific stress response. Lastly, to determine whether heme-binding proteins are required for heme uptake, we incubated hemopexin (Hx) with ZnMP. We observed reduced ZnMP uptake (**Fig. S1e**), indicating that SR-BI specifically mediates the uptake of free heme but not heme-Hx complex. To further characterize how do already known SR-BI ligands affect heme uptake, we conducted a dose-response ZnMP uptake analysis using isolated human HDL at 10, 50, 100 µg/mL. We observed a clear dose dependent inhibition of ZnMP uptake by HDL (**Fig. S1d**). While vector control cells show minimal change at low HDL levels, SR-BI cells exhibit progressive suppression of ZnMP uptake with increasing HDL concentrations.

### Hepatic SR-BI promotes heme clearance and alleviates heme-induced signaling

We next investigated whether hepatic SR-BI contributes to heme clearance (**Fig. 2a**). Upon second hemin injection, we collected blood at various time points and quantified plasma free hemin levels. Wildtype mice exhibited rapid clearance of injected hemin, with 55%, 17% and only 4% remained in circulation at 15-, 30-and 60-minutes post-injection, indicating rapid and effective hemin clearance in wildtype mice. In contrast, AlbcreSR-BI^fl/fl^ mice exhibited impaired hemin clearance, with 98%, 49% and 21% remained in circulation at the same timepoints (**Fig. 2b**). The free hemin concentrations were decreased to baseline levels in wildtype mice but maintained at highly toxic levels (170 µM) in AlbcreSR-BI^fl/fl^ mice at 60-minutes post-injection (**Fig. 2c)**. AlbcreSR-BI^fl/fl^ mice displaced significant accumulation of free hemin in the lung and kidney compared to controls (**Fig. 2d-e**). We also measured free hemin level in the liver. We did not detect a significant difference (**Fig. S2a**). While hepatic SR-BI KO mice lack the cellular entrance of free heme and are expected to show less free heme at tissue level, the highly elevated circulating free heme may account for free heme diffusion into the hepatic tissue. Histologic sections of the kidneys showed stronger eosinophilic staining at the medullary rays in AlbcreSR-BI^fl/fl^ mice (**Fig. 2g**). At higher magnification, this strong eosinophilic staining was identified as interstitial red blood cells (**Fig. 2f, h**, yellow arrow), suggesting damage to the glomerular capillaries, which allowed RBCs to leak into the interstitial space. AlbcreSR-BI^fl/fl^ mice also showed extensive intratubular casts in the medulla and the deposition of pale eosinophilic plugs in the collecting ducts (**Fig. 2h**, fuchsia red arrows), suggesting tubular cell death. Furthermore, we observed clot formation (**Fig. 2g**, turquoise arrow) in AlbcreSR-BI^fl/fl^ mice. These demonstrate that a lack of hepatic SR-BI impairs hemin clearance, resulting in accumulation of free hemin in circulation and organs, which leads to organ injury.

**Fig. 2.**
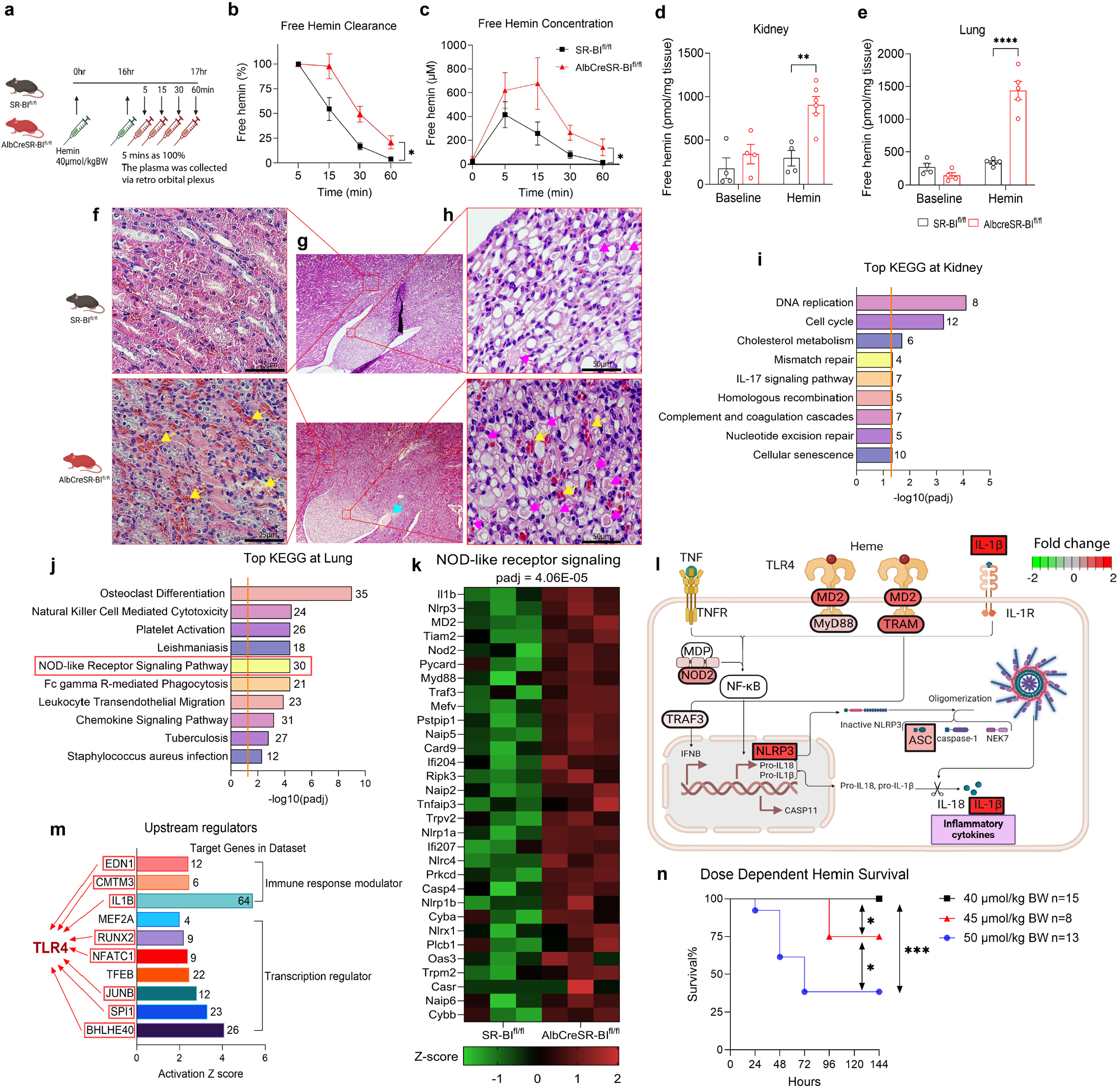
Hepatic SR-BI promotes heme clearance and alleviates heme-induced signaling. (**a**) Experimental design of hemin clearance. SR-BI^fl/fl^ and AlbcreSR-BI^fl/fl^ (n = 4-6) mice were injected with 40 µmol/kg BW of hemin (i.p.). Sixteen hours later, the same dose of hemin was injected (i.v., green syringe) and retroorbital blood was collected at indicated time points for measurement of free heme (red syringe). (**b-c**) Free hemin clearance. The plasma free hemin levels at 5 mins were set as 100% of injected hemin. (**d-e**) Free hemin levels in the kidney and lung tissue lysates were measured one hour post the 2^nd^ hemin injection. (**f-h**) Representative histological images of kidney sections from SR-BI^fl/fl^ and AlbcreSR-BI^fl/fl^ 14 hours post the 2nd hemin injection. (**i-j**) Top KEGG enrichment pathway at kidney and lung. Number next to each bar represents the number of DEGs in that pathway (**k**) Heatmap represents DEGs in NOD-like receptor signaling KEGG pathway. (**l**) Illustrative cartoon of TLR4 and inflammasome activations. (**m**) IPA-predicted upstream regulators. The number within each bar describes target genes from dataset. Activation Z-score ≥ 2 indicates significance. (**n**) SR-BI^fl/fl^ mice were intraperitoneally injected with 5 shots of hemin of various doses. Survival were monitored for 144 hours and analyzed by Log-Rank test; Significances were determined by mixed-effects linear regression and two-way ANOVA. * *P* < 0.05; ** *P* < 0.01; **** *P* < 0.0001.

Free heme is highly active, causing DNA damage through oxidative modification of DNA and organ injury through inflammatory signaling^7^. To collect molecular evidence demonstrating the deleterious consequences of impaired heme clearance, we conducted RNA sequencing (RNAseq). In the kidney, the Kyoto Encyclopedia of Genes and Genomes (KEGG) pathway enrichment analyses identified DNA damage response (including DNA replication, cell cycle, mismatch repair, homologous recombination, nucleotide excision repair, cellular senescence), IL-17 signaling and coagulation as significantly enriched pathways in AlbcreSR-BI^fl/fl^ mice (**Fig. 2i**). These gene profile changes are consistent with the histological evidence of capillary and tubular damage, as well as clot formation, observed in these mice (**Fig. 2f-h**). The activation of these signaling pathways were also reported in high-dose hemoglobin-induced acute kidney injuries in C57BL/6J mice^31^. This similarity suggests that activation of the signaling pathways observed in AlbcreSR-BI^fl/fl^ mice are due to impaired heme clearance, which contributes to kidney injuries.

Several early studies showed that heme induces acute chest syndrome, vaso-occlusive crises and lethality through TLR4-inflammasome activation^32–34^. In the lung, the KEGG pathway enrichment analysis identified NOD-like receptor signaling as a top enriched pathway (**Fig. 2j**). NOD-like receptors are intracellular pattern recognition receptors that detect damage-associated molecular pattern (DAMP) such as heme^35^. Genes such as NLRP3, ASC, and IL-1b, were upregulated 1-2 folds, supporting NLRP3 inflammasome activation. During hemolysis, NOD-like receptors can also act synergistically with Toll-like receptor 4 (TLR4) to illicit potent inflammatory responses. Supporting this synergistic activation, genes involved in TLR4 signaling, including MD2, MyD88, and TRAM, were also upregulated 1-2 folds (**Fig. 2k, l**). We then conducted Ingenuity Pathway Analysis (IPA) to identify upstream regulators. The analysis identified 3 immune response modulators and 7 transcription regulators. Notably, all 3 immune response modulators (EDN1, CMTM3, IL1B) have connections with TLR4 signaling. Among the transcription regulators, 5 out of 7 (RUNX2, NFATC2, JUNB, SPI1, BHLHE40) also show such connections, suggesting that TLR4 drives the differentially expressed genes (DEGs) in the lungs of AlbcreSR-BI^fl/fl^ mice after hemin treatment (**Fig. 2m**). Thus, the RNAseq analysis suggests that the impaired heme clearance induces TLR4-inflammasome activation.

We also looked at whether hepatic SR-BI deficiency might have affected Hb-Hp-CD163 and Hp-Hx-CD91 system, as such change could confound the dependency on SR-BI. We examined the hepatic gene profiles in SR-BI^fl/fl^ and AlbcreSR-BI^fl/fl^ mice and found no changes in these gene expression both at physiological conditions and after hemin treatment (**Fig. S2b**,**c**). At protein level, we examined hepatic CD91 and plasma Hx at baseline and following hemin challenge of early (17hrs) and late crises (38hrs) in both wild-type and SR-BI knockout mice. Both CD91 and Hx level did not differ significantly between two genotypes, ruling out a possible deficiency of the canonical heme scavenging pathway in the SR-BI deficient mice, which may account for their heightened susceptibility towards hemin-mortality (**Fig. S2d**,**e**).

Lastly, we conducted a survival analysis to illustrate that the body is highly sensitive to heme toxicity. We administered various doses of heme to SR-BI^fl/fl^ mice. As shown in (**Fig. 2n**), a slight increase in heme dose from 40 to 45 µmol/kg BW significantly reduced survival from non-lethal to 75%. A dose of 50 µmol/kg BW decreased the survival rate to 38%. This disproportionate rise in lethality underscores the extreme toxicity of free heme, emphasizing the critical importance of efficient free heme clearance in cases of severe hemolysis.

### SR-BI protects against hemolysis- and hemin-induced lethality

We then sought to evaluate the importance of SR-BI as a heme scavenger in physiological conditions and in hemolysis. At physiological conditions, SR-BI^fl/fl^ and AlbcreSR-BI^fl/fl^ mice displayed similar plasma free hemin concentrations and CBC (complete blood count) parameters, including red blood cell (RBC), Hb (hemoglobin), HCT (hematocrit)%, and RET (reticulocyte)% (**Fig. 3a-e**). SR-BI^fl/fl^ and AlbcreSR-BI^fl/fl^ mice also displayed similar spleen weight (**Fig. 3f**). As mentioned above, the RNAseq analysis did not show any difference in gene expression related to heme metabolism (**Fig. S2b-e**). These indicate that a deficiency in hepatic SR-BI does not affect heme levels and RBC homeostasis at physiological conditions.

**Fig. 3.**
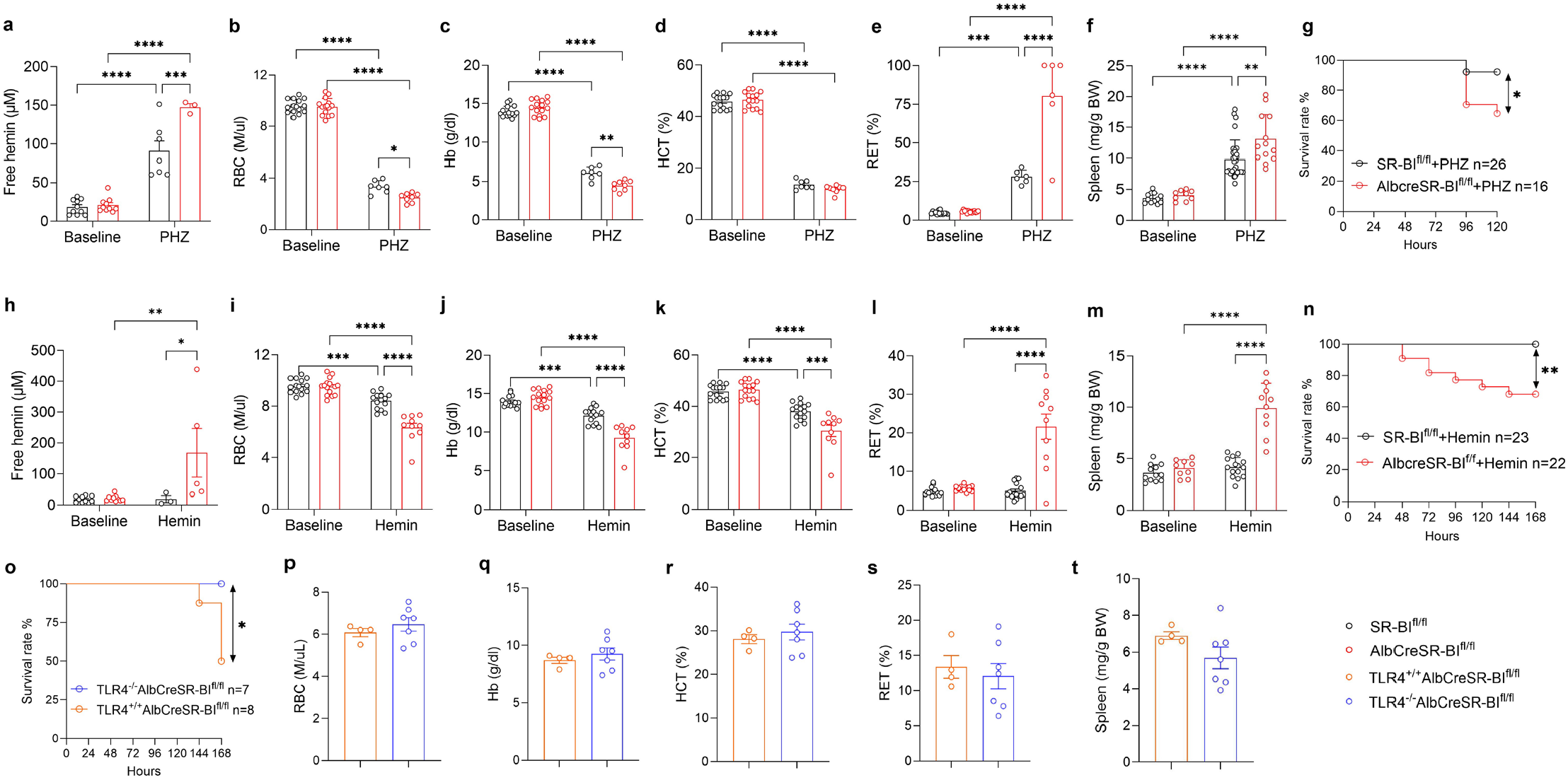
SR-BI protects against hemolysis- and heme-induced lethality. SR-BI^fl/fl^ and AlbcreSR-BI^fl/fl^ mice were intraperitoneally injected with PHZ 30mg/kg BW (4 shots). (**a**) Free hemin levels were measured before and 72 hours after PHZ injections. (**b-e**) The complete blood counts including red blood cells (RBC), hemoglobin (Hb), hematocrit (HCT), and reticulocyte percentages (RET%) were measured at baseline and 120 hrs. SR-BI^fl/fl^ (n= 7-15), AlbcreSR-BI^fl/fl^ (n= 6-15). (**f**) Spleen weights, before and after PHZ injections. SR-BI^fl/fl^ (n=9-30), AlbcreSR-BI^fl/fl^ (n= 9-13). (**g**) Survival rates after PHZ injection. (**h**) SR-BI^fl/fl^ (n=3-10), AlbcreSR-BI^fl/fl^ (n=5-10) mice were intraperitoneally injected with hemin 40 µmol/kg BW (2 shots). Free hemin levels were measured before and 1hr after second hemin injections. (**i-l**) SR-BI^fl/fl^ and AlbcreSR-BI^fl/fl^ mice were intraperitoneally injected with hemin 40 µmol/kg BW (5 shots). The complete blood counts were measured at baseline and 168 hours. SR-BI^fl/fl^ (n=15), AlbcreSR-BI^fl/fl^ (n= 10-15). (**m**) Spleen weights, before and after hemin injections. SR-BI^fl/fl^ (n=15), AlbcreSR-BI^fl/fl^ (n= 9-10). (**n**) Survival rates after hemin injections. (**o**) Survival rates. TLR4^+/+^AlbcreSR-BI^fl/fl^ and TLR4^−/−^AlbcreSR-BI^fl/fl^ littermates were intraperitoneally injected with hemin 45 µmol/kg BW (5 shots). (**p-t**) CBC and spleen weights were obtained at 168 hours. Means ± SEM are plotted. * *P* < 0.05; ** *P* < 0.01; *** *P* < 0.001; **** *P* < 0.0001; Significances were determined by two-way ANOVA and Log-Rank test.

Upon phenylhydrazine (PHZ) challenge to induce hemolysis, both genotypes showed significant increases in free hemin concentrations and decreases in RBC, Hb, HCT%, indicating PHZ induces severe hemolysis (**Fig. 3a-e**). Compared to SR-BI^fl/fl^ mice, AlbcreSR-BI^fl/fl^ mice displayed significantly higher plasma free hemin concentrations, which were associated with lower RBC and Hb, and higher RET% and larger spleen (**Fig. 3a-f**). Highly elevated hemin levels can damage RBCs^36, 37^, leading to secondary hemolysis, which may account for the lower RBC, Hb and higher RET% in AlbcreSR-BI^fl/fl^ mice. This hemin-induced secondary hemolysis (extracellular hemin crisis) has been previously reported in the SCD mice^33^. Additionally, AlbcreSR-BI^fl/fl^ mice had significantly higher WBC counts, with the most prominent increases in neutrophils and monocytes, indicating stronger inflammation (**Fig. S3a-d**). Importantly, PHZ challenge resulted in a significantly higher lethality in AlbcreSR-BI^fl/fl^ mice than in SR-BI^fl/fl^ mice (**Fig. 3g**). These data support that SR-BI is crucial for protection against severe hemolysis by scavenging free heme.

Upon hemin challenge, AlbcreSR-BI^fl/fl^ mice showed significantly elevated plasma free hemin concentrations (**Fig. 3h**). At 168 hours (day 7), both genotypes showed decreases in RBC, Hb, HCT%, indicating hemin induces hemolysis. However, AlbcreSR-BI^fl/fl^ mice displaced significantly lower RBC and Hb, HCT% and higher RET%, along with larger spleens compared to SR-BI^fl/fl^ mice (**Fig. 3i-m)**, which, as explained above, may be caused by hemin-induced secondary hemolysis.

Additionally, the monocyte counts increased significantly in AlbcreSR-BI^fl/fl^ mice (**Fig. S3g**). Importantly, as shown in **Fig. 3n**, administration of hemin at 40 µmol/kg BW were well-tolerated by SR-BI^fl/fl^ mice, but the same dose caused a 30% lethality in the AlbcreSR-BI^fl/fl^ mice. To confirm if any hepatic and metabolic alterations may exist in AlbcreSR-BI^fl/fl^ mice that may cause heightened susceptibility to hemin, we conducted a biochemistry panel for wild-type and AlbcreSR-BI^fl/fl^ mice at baseline. Our data showed that these markers are at similar levels between wild-type and AlbcreSR-BI^fl/fl^ mice (**Fig. S3i-v**).

The upstream regulator analysis (**Fig. 2m**) predicts TLR4 signaling mediates hemin toxicity in hemin-challenged AlbcreSR-BI^fl/fl^ mice. To test this speculation, we generated TLR4^−/−^AlbcreSR-BI^fl/fl^ double knockout mice. As shown in **Fig. 3o**, TLR4^−/−^AlbcreSR-BI^fl/fl^ mice were significantly more resistant to hemin-induced lethality than TLR4^+/+^AlbcreSR-BI^fl/fl^ littermates, despite that hemin induced similar degree of hemolysis in both genotypes (**Fig. 3p-t**).

Taken together, these data support that hepatic SR-BI is essential for protection against severe hemolysis. It functions as a scavenger of free heme to facilitate the clearance of free heme, resulting in less heme toxicity, at least partly, through alleviating TLR4 signaling.

### Upregulation of hepatic SRBI protects against heme-induced lethality

Leveraging a publicly available liver transcriptomic dataset from a humanized SCD mice model (Townes SCD mice) conducted by Grunenwald et al, we analyzed normalized expression counts and heatmap profiles under baseline and heme-induced crisis conditions^38^. Among the hemolysis-scavengers genes examined (SR-BI, CD91, CD163, Hx, Hp), SR-BI displayed the most pronounced trend toward downregulation during crisis, accompanied by the lowest adjusted p-value (**Fig. S4a**). In contrast, canonical scavenger receptors remained comparatively stable. These transcriptomic findings suggest that hepatic SR-BI likely plays a distinct and responsive role in heme clearance during acute hemolytic stress. Their independent study also mirrors our observations in the C57B6J wild-type mice during initial heme crisis. Although two-way ANOVA across baseline, hemin 17hr (early crisis), and hemin 38hr (late crisis) groups did not detect significant changes in SR-BI protein expression, a focused Mann–Whitney test comparing baseline and early crisis revealed a statistically significant downregulation (**Fig. S4b**). Together, this data suggests that SR-BI responds acutely to heme exposure, even if broader time-course comparisons dilute the signal.

Next, we aimed to test the translational potential to target this newly identified heme scavenger. A recent report showed that imatinib, an FDA approved cancer drug, can upregulate hepatic SR-BI^39^. We pretreated mice with imatinib and quantified SR-BI expression in the liver. As shown in (**Fig. 4a-b**), imatinib treatment increased hepatic SR-BI expression by 40%. Then we challenged the SR-BI^fl/fl^ mice with lethal doses of hemin. We observed that imatinib provided significant protection against heme-induced lethality (**Fig. 4c**). Considering that imatinib has functions other than upregulating SR-BI^40^, we aimed to validate the rescue dependency on SR-BI mechanism. First, we tested imatinib efficacy in AlbCreSR-BI^fl/fl^ mice. We found that imatinib no longer conferred a significant survival benefit, suggesting that SR-BI is a major mediator of its protective action against hemin-mortality (**Fig.S4c**). Next, to confirm the protection is specifically mediated by SR-BI, we used adenovirus overexpression approach. To further enhance translation from mice findings to human physiology, we used adenovirus containing human SR-BI sequence (Ad.hSR-BI). Injection of

**Fig. 4.**
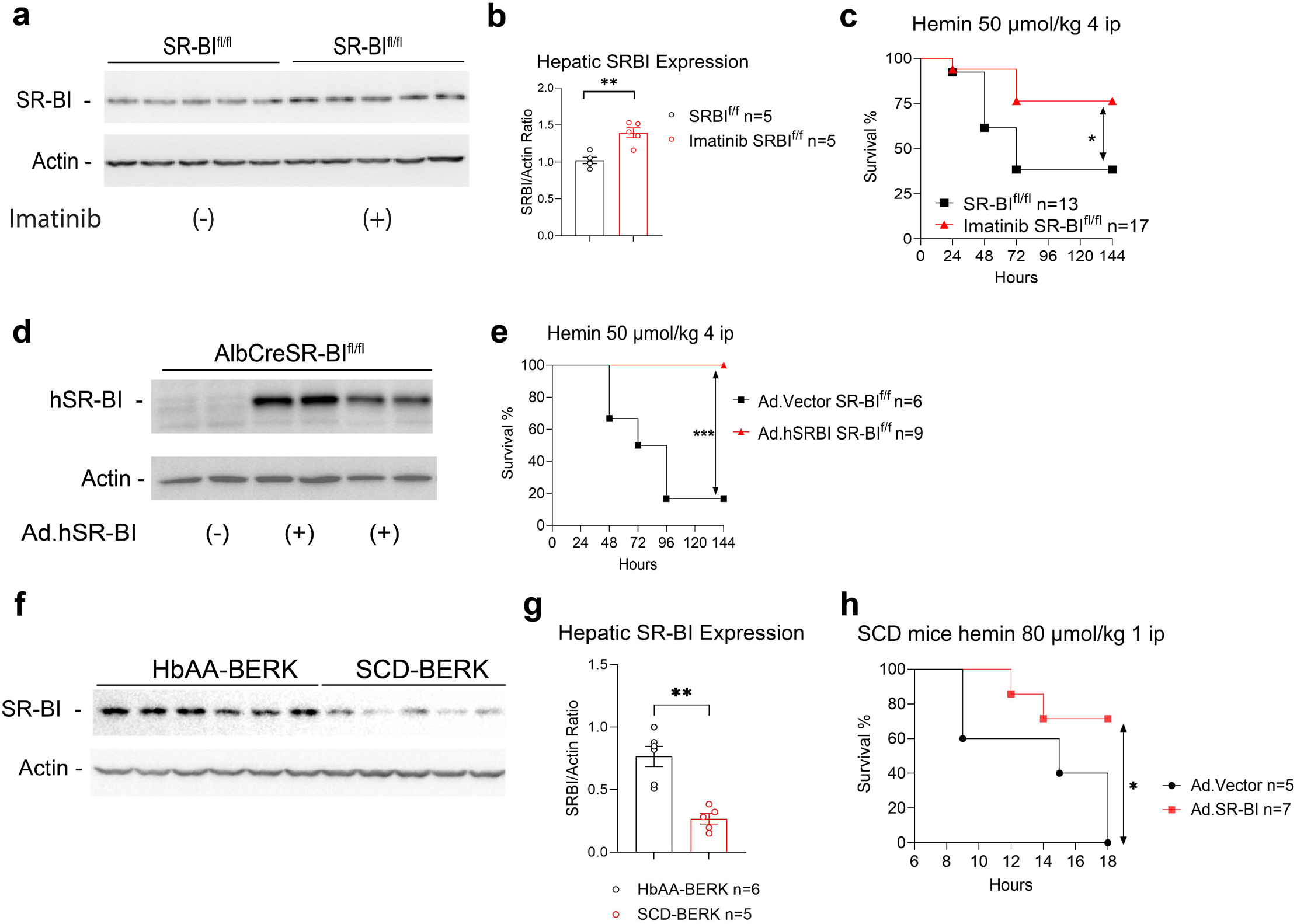
Upregulation of hepatic SRBI protects against heme-induced lethality. (**a-b**) Hepatic SR-BI expression in SR-BI^fl/fl^ mice treated with/without imatinib (50 mg/kg BW) for 3 days, followed by daily hemin injection of 40 μmol/kg for 5 days. (**c**) Survival analysis of SR-BI^fl/fl^ mice treated with/without imatinib for 3 days, followed by daily hemin injection. (**d**) Hepatic SR-BI expression in AlbcreSR-BI^fl/fl^ mice treated with adenovirus expressing human SR-BI (Ad.hSR-BI) or vector (Ad.vector) for 3 days. (**e**) Survival analysis of SR-BI^fl/fl^ mice treated with Ad.hSR-BI or Ad.vector for 3 days, followed by daily hemin injection. (**f-g**) Hepatic SR-BI expression in Non-SCD mice (Hba^−/−^Hbb^+/−^Tg^+/−^ n=6) and SCD-BERK littermates (Hba^−/−^Hbb^−/−^Tg^+/−^) after daily hemin injection of 45 μmol/kg for 5 days. (**h**) Survival analysis of SCD-BERK mice treated with Ad.SR-BI or Ad.vector for 3 days, followed by hemin injection. Data are shown as means ± SEM. * *P* < 0.05; ** *P* < 0.01; *** *P* < 0.001, determined by Student’s t test. Survival curves were analyzed by the Log-Rank test.

Ad.hSR-BI induced substantial hSR-BI expression in the liver (**Fig. 4d**). Then we pretreated SR-BI^fl/fl^ mice with either Ad.vector or Ad.hSR-BI and challenged them with hemin. The overexpression of hSR-BI offered significant protection against heme-induced lethality (**Fig. 4e**). Additionally, no significant untoward effect of Ad.Vector was observed (**Fig.S4d**). These results provide a proof-of-concept that hepatic SR-BI is a potential therapeutic target for hemolysis. To better understand how SR-BI integrates with existing heme detoxification systems, we evaluated if Hp and Hx levels are altered following therapeutic SR-BI upregulation. Immunoblots show that Hp and Hx levels remain largely unchanged, suggesting that hepatic SR-BI operates as an independent heme uptake mechanism (**Fig. S4e**).

Given liver SR-BI’s critical importance during hemolysis and considering that liver cirrhosis is common in SCD patients^41^, we asked whether SR-BI expression would be affected by sickle cell hepatopathy. We examined hepatic SR-BI expression in the Berkeley mice (SCD-BERK), a more severe form of humanized SCD model. These mice express human sickle hemoglobin and display liver infarcts and injuries similar to those seen in humans^42^. The immunoblot analysis revealed a significant reduction of more than 50% SR-BI expression in SCD mice compared to their non-SCD HbAA littermates (**Fig. 4f-g**). Given the substantial reduction in SR-BI expression, and prior studies demonstrating that imatinib has beneficial effects in SCD mice^43, 44^ and in SCD patients^45–48^, we next sought to determine whether imatinib may upregulate hepatic SR-BI in the context of SCD. The immunoblot analysis showed a significant upregulation of hepatic SR-BI in SCD mice after imatinib treatment (**Fig. S4f**). To mechanistically test if upregulating SR-BI levels could rescue hemin toxicity, we used an adenoviral approach to increase liver SR-BI in these SCD mice and subsequently challenged them with a lethal dose of hemin. Notably, SR-BI gain-of-function offered significant protection against hemin-induced death (**Fig. 4h**). These findings, demonstrated in the humanized SCD mice model, consistently support that SR-BI has a strong therapeutic potential in the context of SCD.

## Discussion

In this study, we employed multiple approaches to identify hepatic SR-BI as a scavenger of free heme that is essential for protection against severe hemolysis. Using computer modeling and SPR, we showed that SR-BI binds heme at pathological concentrations of heme; using flow cytometry and confocal analysis, we showed that SR-BI facilitates the uptake of free heme in cells stably expressing SR-BI and in primary hepatocytes; using HO-1 expression as a marker of intracellular heme uptake, we confirmed that SR-BI facilitates heme uptake. Using hepatic SR-BI null mice (AlbcreSR-BI^fl/fl^), we demonstrated that hepatic SR-BI facilitates heme clearance, protects heme-induced organ injury and lethality. By enhancing hepatic SR-BI expression with imatinib and Ad.hSR-BI, we provided a proof- of-concept that hepatic SR-BI is a potential therapeutic target for severe hemolysis. Finally, we showed reduction in hepatic SR-BI expression in humanized SCD mice, suggesting that the loss of hepatic SR-BI might be a risk factor for SCD crises. Importantly, upregulating hepatic SR-BI expression rescued the SCD mice. The present study identifies hepatic SR-BI as a targetable scavenger of free heme in hemolysis. (**Fig. 5**).

**Fig. 5.**
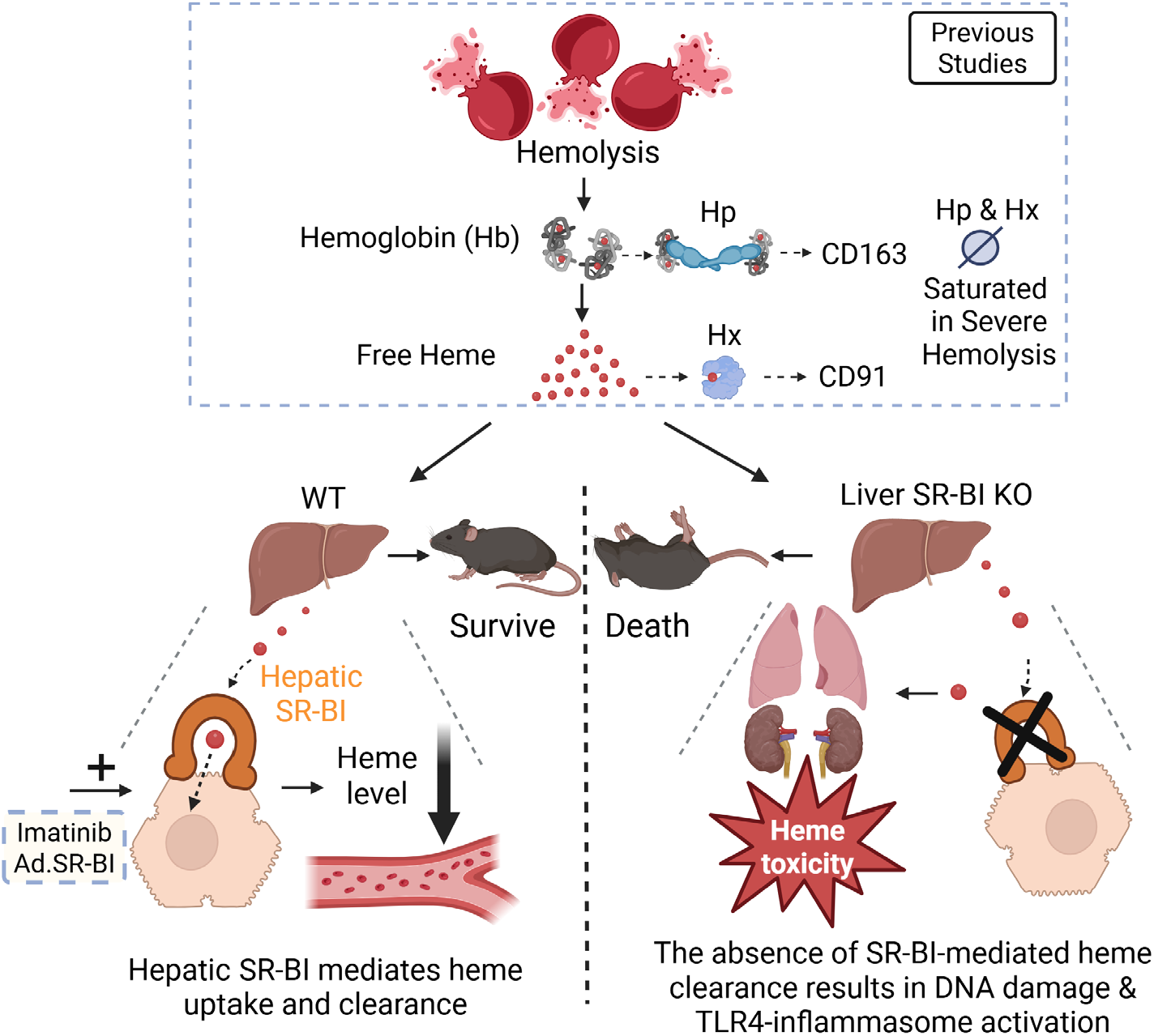
Schematic model of hepatic SR-BI as a heme scavenger that is essential for protection against hemolysis. During hemolysis, hemoglobin (Hb) and heme released from red blood cells are bound by haptoglobin (Hp) and hemopexin (Hx), respectively. These complexes are then removed by CD163 and CD91 cells. However, in severe hemolysis, these systems become saturated due to the depletion of Hp and Hx, leading to dangerous levels of free heme. Scavenger receptor BI (SR-BI) binds heme when plasma heme reaches pathological levels. In the absence of liver SR-BI, free heme remains highly elevated in the plasma, triggering potent inflammatory and DNA damage responses in the lung and kidney respectively. This leads to a heightened lethality. Conversely, in the presence of hepatic SR-BI, plasma heme is rapidly cleared, protecting against heme-induced injury and mortality. Upregulating hepatic SR-BI via imatinib or adenoviral delivery protects wild-type and SCD mice against heme toxicity.

To date, no crystal structures have been published for SR-BI, however, with the advent of diffusion-based generative models, high quality protein structure predictions can be readily accessed via databases such as UniProt. In the present study, we leveraged high-confidence AlphaFold3 structural predictions and protein–protein docking to model the SR-BI dimer, the functional unit required for efficient lipid transport^49^. A protein-protein dimer pose was selected that showed similarity to that of the previously published work of Koenig et al., showing that the G319V mutation provided additional stabilizing interactions between the monomers of SR-BI, with interactions between the CPC motif of one dimer and the and the helix-turn-helix motif of residues 152-166^50^. To experimentally probe the binding interface in lieu of these structural methods, we examined the well-characterized SR-BI C323G mutant, which is correctly trafficked to the plasma membrane yet lacks HDL-derived cholesterol ester uptake^27^. The C323G mutation shows a notable loss in hemin uptake and is primarily caused by two overall factors. When mutated, the C323G mutation breaks the disulfide bond located at the CPC (Residues 321 to 323) motif, a critical recognition motif that is conserved across the CD36 family of proteins, this breakage leaves a free cysteine residue to form errant dimers with other mutated versions of SR-BI, leading to malformation of the dimer structure and loss of protein functionality^28, 29, 51^. Even when the correct dimer positioning is able to form, the breakage of the disulfide bond leads to a contraction of the binding site, causing the central proline of the CPC motif to descend down into the binding site that Hemin occupies, blocking the WT binding mode of Hemin. In SR-BI C323G mutant, residues Y194 and T333 further contract the SR-BI binding site, destroy the strong hydrogen bond with N330, clashing against the peripheral methyl and vinyl carbons of the protoporphyrin ring. Additionally, the energy minimization of the C323G SR-BI dimer shows the K151 residue repositioning away from the center of the binding site, increasing the distance between it and Hemin by over 1.5 Å, reducing the favorable coulombic interactions between the two. Additionally, the cleavage of the C321–C323 disulfide bond results in an additional free thiol, decreasing the net positive charge of SR-B1 and thus reducing the overall coulombic attraction between SR-BI and Hemin. These clashes within the binding site along with the malformed dimer creation, are responsible for the loss in Hemin transport by SR-BI.

Hemolysis occurs under both physiological and pathological conditions. Under normal physiology, approximately 20-40 billion RBCs (equivalent to 0.6 to 1.2 grams of Hb) undergo hemolysis every day, which releases up to 74 µmoles of free heme into circulation^6, 52^. With dissociation constants (Kd) in the pico- to nanomolar range, Hp and Hx can efficiently sequester free Hb and heme, which keeps free Hb and heme at extremely low levels (0.2 µM) under physiological conditions^32^. HpHx double knockout mice have 1.7-fold larger spleens than wildtype mice^9^, suggesting their critical protective roles at physiological conditions. However, during pathological hemolysis, such as SCD crisis, malaria, and sepsis, both Hp and Hx are often depleted, limiting their protective capacity. This speculation is supported by an earlier study that HpHx double knockout mice paradoxically have higher survival rates compared to wildtype mice upon severe hemolysis challenge^9^.

In contrast to high affinity binding between heme and heme binding proteins, SR-BI binds to heme with low affinity with a Kd at 18.2 µM. Given the average free heme levels in SCD and malaria patients at 70 µM ^21, 22^, SR-BI may effectively scavenge heme at pathological conditions and confer protection. Indeed, heme clearance experiment showed that the wildtype mice rapidly and effectively cleared injected hemin to baseline levels within 60 min but the free hemin were still at highly toxic levels (170 µM) at 60 min post injection in the AlbcreSR-BI^fl/fl^ mice. Importantly, AlbcreSR-BI^fl/fl^ mice were significantly more susceptible to PHZ-induced hemolysis and hemin-induced toxicity. Considering that there is no difference in RBC, Hb, RET and spleen weight at physiological conditions between SR-BI^fl/fl^ AlbcreSR-BI^fl/fl^ mice, our study supports hepatic SR-BI as an efficient scavenger of free heme in pathological hemolysis.

There are some differences in hemolysis scavenging systems between rodents and humans. For examples, Hx is an acute phase protein in rodents but not in humans^53^. Additionally, mouse CD163 binds to both the Hb-Hp complex and free Hb, however human CD163 only recognizes the Hb-Hp complex^3^. In contrast to these differences, SR-BI biology appears highly conserved between rodent and human: 1) both SR-BI null mice and SR-BI mutations in humans show impaired HDL metabolism, decreased adrenal stress response and increased risk of cardiovascular disease^14^; 2) the SR-BI extracellular domain shares 80% sequence identity between rodents and humans^54^. Importantly, the amino acids involved in heme binding are either conserved or maintained their hydrophobicity (**Fig. S1**), suggesting that both mouse and human SR-BI would exhibit a similar heme binding affinity; 3) In the present study, we conducted parallel experiments using both mouse and human SR-BI. In vitro, both primary hepatocytes (mouse SR-BI) and CHO cells (human SR-BI) showed dose-dependent heme uptake. In vivo, both endogenous mouse SR-BI overexpression and adenoviral human SR-BI overexpression provided protection against heme toxicity. Thus, this study provides valuable insights into our understanding of protection against heme toxicity in human pathological conditions such as sepsis, SCD and malaria.

SCD patients frequently seek medical care in hospitals due to vaso-occlusive crises. These crises can be complicated by acute organ dysfunction such as stroke or acute kidney injury, creating long-lasting damage. Thus, in addition to crises resolution, a preventive therapeutic approach is needed to avoid irreversible damage associated with SCD crises. Early studies demonstrated the efficacy of using Hx to scavenge heme in SCD crisis resolution^36, 55–61^. Notably, treatment that specifically targets free heme by using plasma-derived human Hx demonstrated excellent safety and tolerability, now entering into a phase 2 clinical trial of SCD. These studies not only indicate that free heme is a promising drug target but show exciting progress for an unmet need of SCD crises treatment.

Building on these prior elegant studies, leveraging liver SR-BI to boost free heme scavenging introduces a novel strategy to target heme toxicity and could serve as a preventative approach. In this study, imatinib pretreatment significantly increased SR-BI expression in the liver and prevented heme toxicity. Previous studies have shown that imatinib protects against sickle-cell related injury in SCD mice^43, 44^. Several case reports and a recent clinical trial showed that imatinib reduced pain crises in patients with SCD^45, 48^. Our study provides a novel mechanistic insight by which imatinib protects against heme toxicity, a critical contributor of pain crises. Additionally, adenoviral overexpression of human SR-BI reduced hemin-induced lethality in the wildtype mice. These results offer a proof-of-concept that inducing liver SR-BI expression may serve as a preventative measure against frequent heme insults observed in SCD. Lastly, we demonstrated that sickle cell hepatopathy may significantly reduce hepatic SR-BI expression, which could be a potential risk factor for worse SCD outcomes. Enhancing liver SR-BI expression in SCD mice confers a significant protection against acute heme toxicity. These results consistently support the therapeutic potential of SR-BI against heme toxicity.

While our study identifies SR-BI as a key determinant of cellular heme handling and demonstrates that SR-BI upregulation confers robust protection against acute heme-induced lethality, several important questions remain. SCD is characterized by chronic, low-grade hemolysis, and it is unknown whether SR-BI–targeted strategies provide sustained benefit in this long-term setting. Although curative gene therapy has transformed the therapeutic landscape for SCD, its’ widespread implementation is limited by cost and accessibility^62^. Imatinib, the first-generation BCR-ABL tyrosine kinase inhibitor now available as an inexpensive generic drug, represents a potentially scalable intervention, especially in resource-limited settings. Determining whether chronic imatinib treatment can improve long-term outcomes in SCD may help bridge the gap until curative therapies become broadly accessible.

The uncovering of a novel heme scavenging pathway expands the toolbox for addressing the unmet need for treatment against heme toxicity in SCD crises. Restoring SR-BI expression and promoting heme scavenging via SR-BI might therefore offer a new avenue to treat SCD, a disease affecting approximately 8 million people worldwide. In conclusion, our proof-of-concept experiments highlight the therapeutic potential of hepatic SR-BI in mitigating the harmful effects of free heme in severe hemolysis, which may represent a new therapeutic category in combating hemolytic complications.

## Author contributions

M.I. and J.X. performed most of the studies and wrote the manuscript. L.G, D.H. and Q.W. contributed to experiments. A.H.W. and C.-G.Z. performed in-silico docking experiments. A.J. and P.S. provided murine SR-BI adenovirus. W.S, S.L, Z.G, and M.G. performed histological analysis. G.S. provided isolated human HDL. B.H. provided advice on statistical analysis. J.J provided lysotracker and helped with confocal imaging. C.M. and P.W.S. generated SR-BI^fl/fl^ mice. X.-A.L. conceptualized and oversaw studies, assisted in manuscript preparation, and procured funding.

## Acknowledgments

This study was supported by NIH R35 GM141478 (X.-A.L.); VA 1I01BX004639 (X.-A.L.); NIH R01 HL126795 (C.M.); NIH R01 HL181134 (P.W.S.); American Heart Association AHA-predoc-fellowship 26PRE1563237 (J.X.). Its contents are solely the responsibility of the authors and do not necessarily represent the official views of the NIH, VA or AHA.

## Materials and Methods

### Materials

Hemin (H9039), PHZ (114715) were from Sigma-Aldrich. ZnMP (M40628) was from Frontier Scientific. Other materials are listed in **Major Resources Tables**.

### Mice

SR-BI^fl/fl^ mice were generated as described^15^. The mice were in C57BL/6J background^63^. Albcre mice in C57BL/6J background were from the Jackson Laboratory (Stock No:003574). SR-BI^fl/fl^ and Albcre mice were bred to generate AlbcreSR-BI^fl/fl^ and SR-BI^fl/fl^ littermates. TLR4^−/−^ mice were from the Jackson Laboratory (Stock No:007227). TLR4^−/−^ mice were bred with AlbcreSR-BI^fl/fl^ to generate TLR4^−/−^AlbcreSR-BI^fl/fl^ double knockout mice and TLR4^+/+^AlbcreSR-BI^fl/fl^ littermates. SCD-Berk mice were from Jackson Laboratory (Stock No:003342). 12-20 week-old littermates were randomly allocated into the groups for all animal experiments. Gender-match was considered whenever possible. Blinding was performed for survival and biochemical analyses. No data were excluded. The animals were fed with standard laboratory chow diet. Animal care and experiments were approved by the Institutional Animal Care and Use Committee of the University of Kentucky.

### Adenovirus experiment

The adenovirus of human SR-BI has the human SR-BI expression cassette and E1&E3 self-replication genes deleted. The Ad.vector does not have the human SR-BI expression cassette. Adenovirus of mouse SR-BI were generated as previously described^64^.

### Analysis of hemin clearance

Hx presents in circulation at high levels at physiological conditions but depleted in severe hemolysis. To mimic this condition, Dr. Pawlinski’s group developed a two-heme-injection protocol to deplete circulating Hx by the first heme injection and then determine heme toxicity by the second heme injection^65^. To evaluate free heme clearance in hemolysis, we did two hemin injections followed this protocol. Briefly, SR-BI^fl/fl^ and AlbcreSR-BI^fl/fl^ mice were intraperitoneally preconditioned with hemin at 40 μmol/kg BW to deplete Hx. Sixteen hours later, the mice were intravenously injected with hemin at 40 μmol/kg BW. Then, plasma was collected via retro orbital venous plexus at 5, 15, 30 minutes after hemin injections. The mice were terminated after 60 minutes. Plasma and organs were collected for free hemin quantification using a free Hemin Assay Kit (MAK036, Sigma). The plasma free hemin levels at 5 min were set as 100% of injected hemin and then the clearance of hemin at each time point was determined.

### PHZ-induced hemolysis and hemin-induced toxicity

Phenylhydrazine (PHZ; Sigma) was freshly prepared by dissolving in sterile phosphate-buffered saline (PBS), and the pH was adjusted to 7.4 using 1 M NaOH. Hemin was dissolved in 0.9% saline containing D-sorbitol and sodium carbonate, then filtered through a 0.22 μm membrane, as previously described^32^. To induce hemolysis, age- and sex-matched mice were injected intraperitoneally with PHZ at a dose of 30 mg/kg body weight (BW) every 24 hours for four consecutive days. Mice were euthanized 120 hours after the first injection for blood and tissue collection and analysis. For hemin challenge, mice received intraperitoneal injections of hemin at 40 μmol/kg BW every 24 hours for a total of five doses. Animals were euthanized 168 hours after the initial injection for sample collection. TLR4^−/−^ mice are partly in C57BL/6J background and resistant to hemin-lethality. Thus, a higher hemin dose of 45 μmol/kg BW were used for TLR4^−/−^ AlbcreSR-BI^fl/fl^ double knockout mice and TLR4^+/+^AlbcreSR-BI^fl/fl^ littermates. SCD-Berk mice also exhibited resistance to hemin challenge.

Therefore, a higher dose of 80 μmol/kg BW was used in mice treated with Ad.SR-BI or control vector. Complete blood counts (CBC) were performed using the IDEXX CBC analyzer (IDEXX Laboratories).

### Analysis of cellular uptake of ZnMP by flow cytometry

CHO and HEK293 cells stably expressing vector, SR-BI WT and SR-BI C323G mutant were generated as previously described^66^. The ZnMP uptake was evaluated by flow cytometry as previously described^26^. Briefly, CHO cells expressing either vector or SR-BI were cultured to 70-80 % confluency. The cells were washed with PBS and then incubated with Zn(II) Mesoporphyrin IX (ZnMP). After incubation for 15 minutes at 37°C, unbound ZnMP was removed by adding 2 ml of ice-cold 5% BSA in PBS. The cells were then collected by Trypsin/EDTA on ice to remove cell surface ZnMP, washed with PBS three times. Cell pellets were resuspended in FACS buffer, and the fluorescence intensity of ZnMP was measured using a FACSymphony cell analyzer (BD Biosciences). Flow cytometry data was analyzed using FlowJo software (Treestar). To assess ZnMP uptake, the MFI of the control was set to 100, and the MFI of the treatment groups was converted to relative values with the control expressed as %MFI. Primary hepatocytes from liver tissue samples were prepared as previously described^67^. Viable cells were counted and plated to collagen IV-coated plates at a density of 2 x 10^5^ cells per well. After 6 hours, the cells were washed and the ZnMP uptake assay was performed as described above.

### Analysis of cellular uptake of ZnMP by fluorescence microscopy

Cells were seeded on a Nunc™ Lab-Tek™ II Chamber Slide™ System (Thermo Fisher Scientific) at density of 3.0 x 10^4^ cells in 0.4 ml in Ham’s F-12 medium, and were incubated for 24 hours. 20 µM of ZnMP in Ham’s F-12 was introduced. For competition studies, 100 µM of hemin was added. For inhibition studies, cells were preincubated for 1 hour with BLT1 10 µM and then proceed to ZnMP uptake studies. After 30 minutes of incubation, the ZnMP uptake was stopped by adding 0.5 ml of ice-cold 5% BSA in PBS. The cells were then washed three times with PBS before being mounted with ProLong™ Gold Antifade Mountant with DAPI. Immunofluorescence microscopy was conducted using a confocal A1R system (Nikon), and images were acquired and processed with the Nikon software.

### Surface plasmon resonance (SPR) platform

Fc Tagged human SR-BI and hemin were sent to Creative Biolabs (Shirley, NY, USA) for SPR analysis. The SR-BI was immobilized onto the CM5 sensor chip surface at an immobilization level of approximately 15,000 RU. Hemin at the concentration of 50, 25, 12.5, 6.25, and 3.125 μM was injected into the sensor surface to observe its interaction with SR-BI protein. The 1:1 binding model was used to determine the binding affinity and/or kinetics. Using the 1:1 binding model, the kinetic constants of the hemin binding to SR-BI protein were measured.

### RNA-sequencing analysis

SR-BI^fl/fl^ and AlbcreSR-BI^fl/fl^ mice were treated with/without hemin at 40 µmol/kg BW, i.p. Twenty-four hours later, a second injection was performed. Thirty-eight hours post the initial injection, RNA was isolated from Kidney, lung and liver, and sent to Novogene for RNA sequencing analysis as recently described^68, 69^. We used n=3, considering the robust differences between SR-BI^fl/fl^ and AlbcreSR-BI^fl/fl^ mice. To compensate for the small sample size, we used deep sequencing depth and paired-end sequencing with a read length of 150 base pairs. Additionally, we applied stringent statistical thresholds of FDR <0.05 and |Log2(Foldchange)| ≥ 1 when looking at gene individually. This approach allowed us to achieve goal to prioritize dramatic changes over detecting subtle differences^68, 69^. Ingenuity Pathway Analysis (IPA) was used to analyze the gene expression patterns.

### Immunoblotting

The assay was conducted as previous described^66^.

#### Statistical Analysis

The data were analyzed using GraphPad Prism 10. The survival assay was analyzed using Log-Rank test and Kaplan-Meier plots. For experiments comparing two groups, significance was determined using a two-tailed Student t-test. For experiment comparing more than two groups, significance was evaluated by One-Way or Two-Way ANOVA, followed by post hoc analysis using Tukey’s test. The data were means ± SEM.

## Data Availability

The datasets generated and/or analyzed during the current study are available from the corresponding author on request. All raw data supporting the findings of this study have been deposited in Figshare DOI (https://doi.org/10.6084/m9.figshare.30089977). Source data for figures and tables are provided with the paper.

