## Supplemental Figures for "Identification of scavenger receptor BI as a scavenger of free heme that is essential for protection against hemolysis"

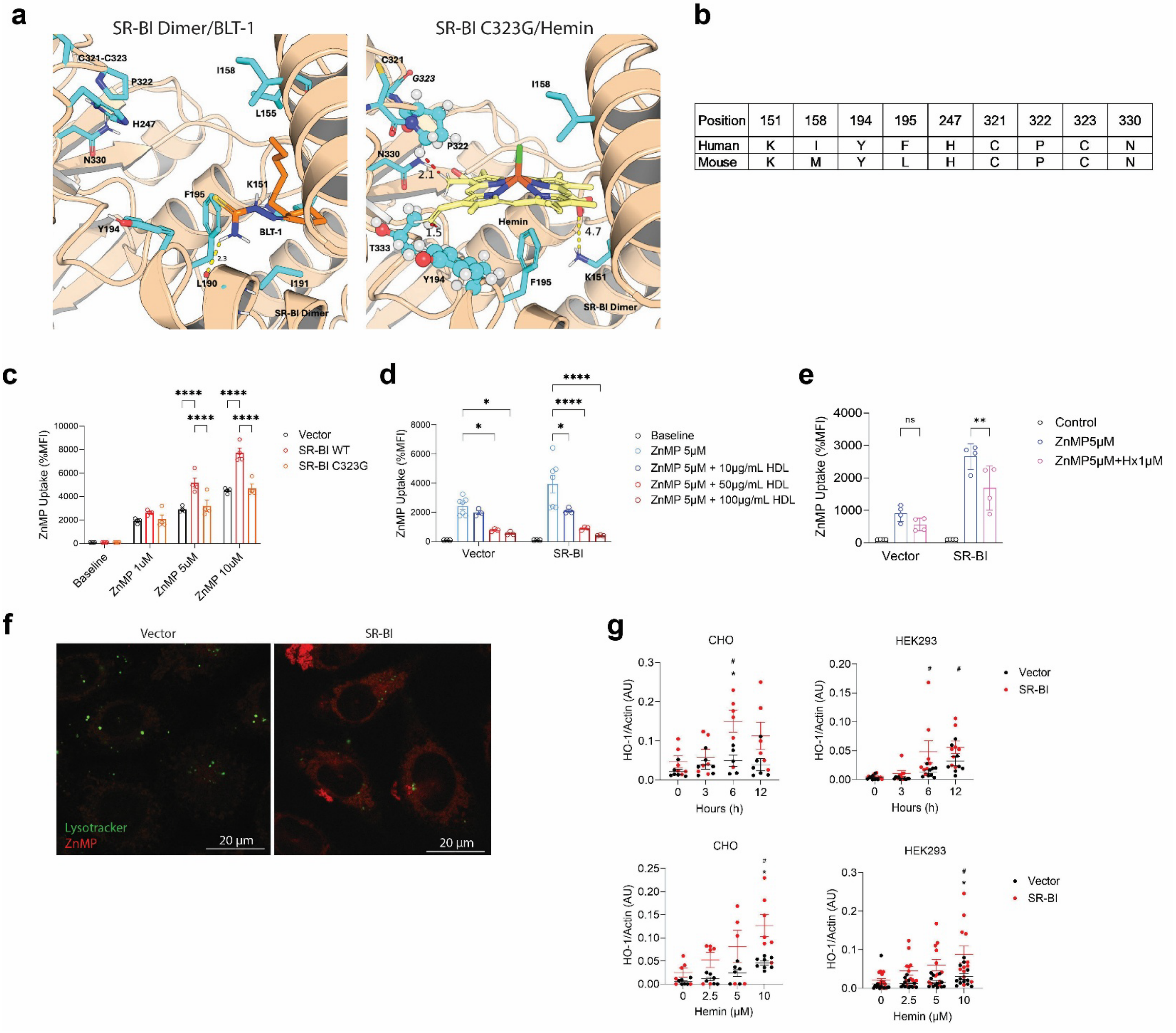


| Position | 151 | 158 | 194 | 195 | 247 | 321 | 322 | 323 | 330 |
|---|---|---|---|---|---|---|---|---|---|
| Human | K | I | Y | F | H | C | P | C | N |
| Mouse | K | M | Y | L | H | C | P | C | N |

**Fig. S1. SR-BI facilitates the uptake of free heme.**

(**a**) Binding modes of SR-BI WT with BLT-1, and SR-BI C323G mutant with hemin, predicted using an energy-minimization workflow implemented in AutoDock Vina. (**b**) Amino acid sequences similarity between human and mouse SR-BI involved in hemin binding. (**c**) Flow cytometry analysis of ZnMP uptake in CHO cells expressing vector, SR-BI WT or SR-BI C323G mutant. (**d**) Flow cytometry analysis of ZnMP uptake in CHO cells expressing vector and SR-BI WT, in the presence of 10, 50, 100 µg/ml high-density lipoprotein (HDL) and (**e**) in the presence of 1 µM hemopexin (Hx). (**f**) CHO cells expressing vector or SR-BI were incubated with 20 µM ZnMP for 30mins and co-stained with 500 nM LysoTracker for 1hr to assess intracellular localization. (**g**) Individual data point for hemin time and dose-dependent HO-1 expression in Figure 1 (**m-p**). The * denotes comparison with the vector control at the same timepoint, while # denotes comparison with baseline. Means ± SEM are plotted from three independent experiments. * $P < 0.05$; ** $P < 0.01$; *** $P < 0.001$; **** $P < 0.0001$; Significances were determined by two-way ANOVA.

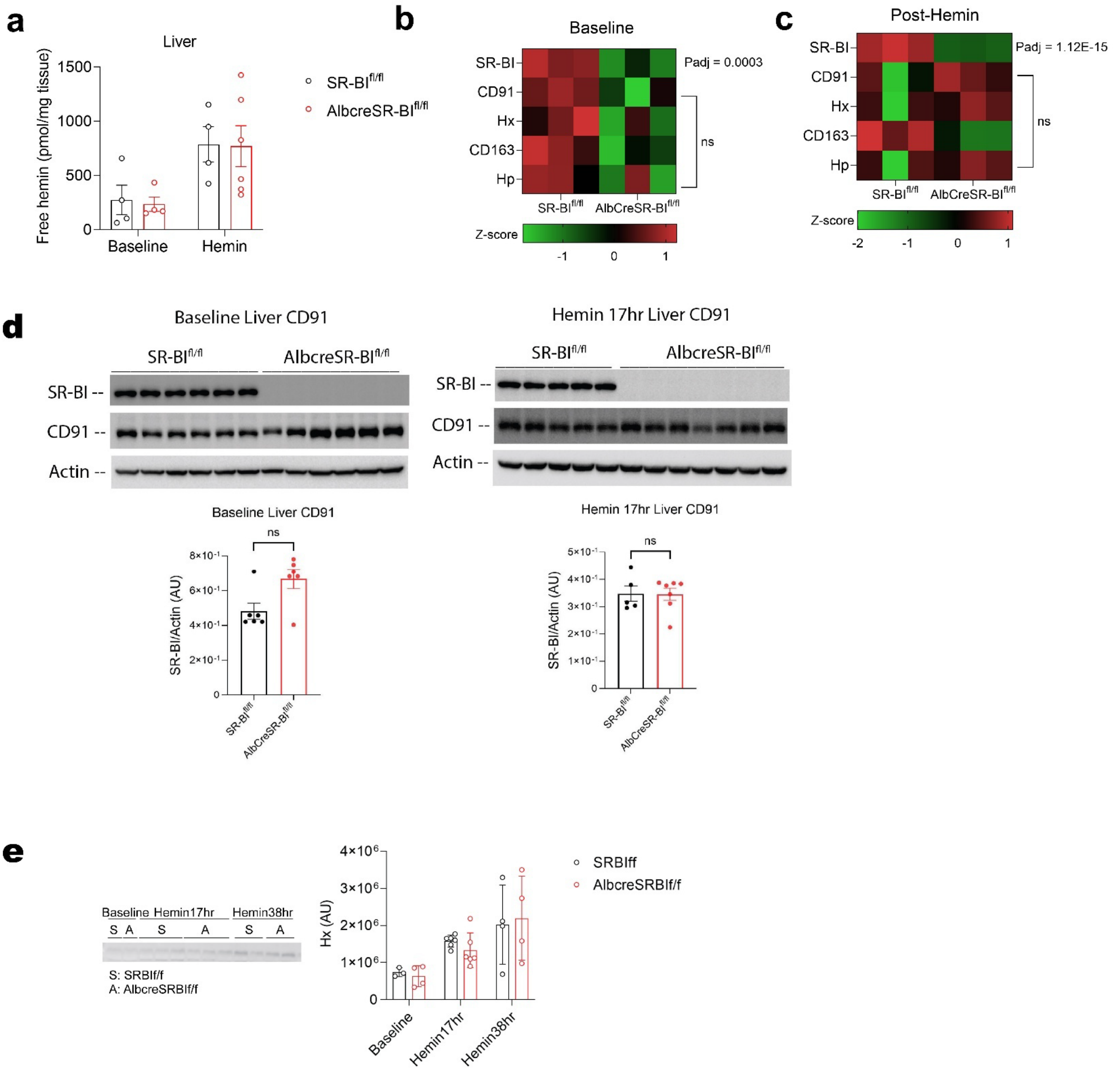


**Fig. S2. Liver gene profiles of SR-BI$^{fl/fl}$ and AlbcreSR-BI$^{fl/fl}$ mice.**

(**a**) Free hemin levels in the liver tissue lysates were measured one hour post the 2$^{nd}$ hemin injection from the free hemin clearance study. (**b-c**) Heatmap of genes of interests from SR-BI$^{fl/fl}$ and AlbcreSR-BI$^{fl/fl}$ mice liver at baseline and after hemin treatment. None significant (ns) is based on adjusted p-value. (**d-e**) SR-BI$^{fl/fl}$ and AlbcreSR-BI$^{fl/fl}$ ($n = 3$-$6$) mice at baseline and after one injection of 40 µmol/kg BW of hemin (i.p.). 30 µg of liver lysate and plasma protein were loaded and examined for SR-BI, CD91, Actin, and Hx via immunoblot. Significance was determined by two-way ANOVA and Mann-Whitney test.

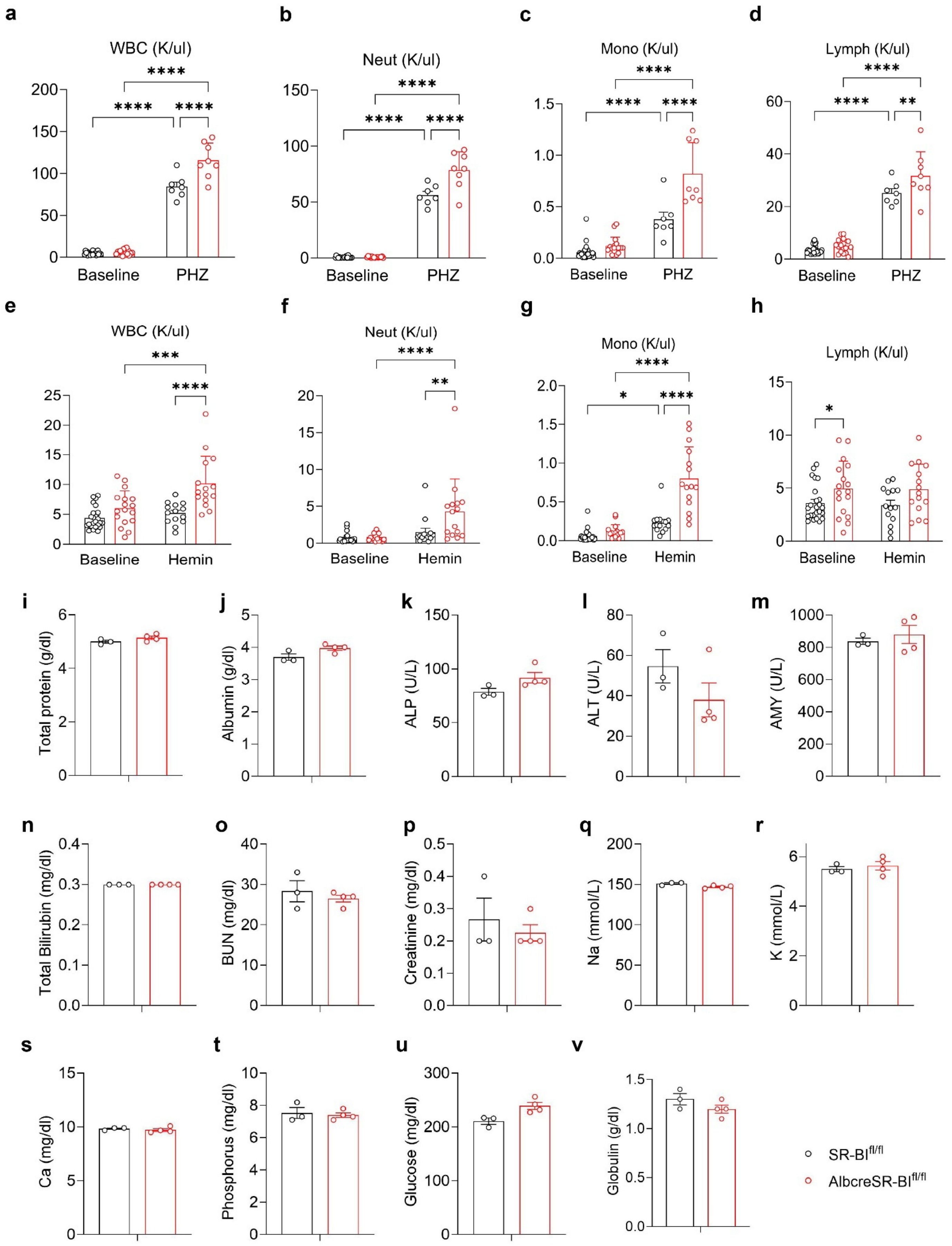
a
WBC (K/ul)
b
Neut (K/ul)
c
Mono (K/ul)
d
Lymph (K/ul)
Baseline
PHZ
e
WBC (K/ul)
f
Neut (K/ul)
g
Mono (K/ul)
h
Lymph (K/ul)
Baseline
Hemin
i
Total protein (g/dl)
j
Albumin (g/dl)
k
ALP (U/L)
l
ALT (U/L)
m
AMY (U/L)
n
Total Bilirubin (mg/dl)
o
BUN (mg/dl)
p
Creatinine (mg/dl)
q
Na (mmol/L)
r
K (mmol/L)
s
Ca (mg/dl)
t
Phosphorus (mg/dl)
u
Glucose (mg/dl)
v
Globulin (g/dl)
SR-BI$^{fl/fl}$
AlbcreSR-BI$^{fl/fl}$

**Fig. S3. CBC in mice treated with PHZ and hemin. Baseline biochemistry panel**

Total white blood cells (WBC), neutrophil (Neut), monocytes (Mono), lymphocytes (Lymph) in SR-BI$^{fl/fl}$ and AlbcreSR-BI$^{fl/fl}$ mice after intraperitoneal injections with PHZ 30 mg/kg BW (4 shots) (**a-d**) and hemin 40 µmol/kg BW (5 shots) (**e-h**). (**i-v**) Biochemistry panel of SR-BI$^{fl/fl}$ and AlbcreSR-BI$^{fl/fl}$ at baseline. Means ± SEM are plotted. * $P < 0.05$; ** $P < 0.01$; *** $P < 0.001$; **** $P < 0.0001$; Significances were determined by two-way ANOVA for (**a-h**) and Mann-Whitney for (**i-v**).

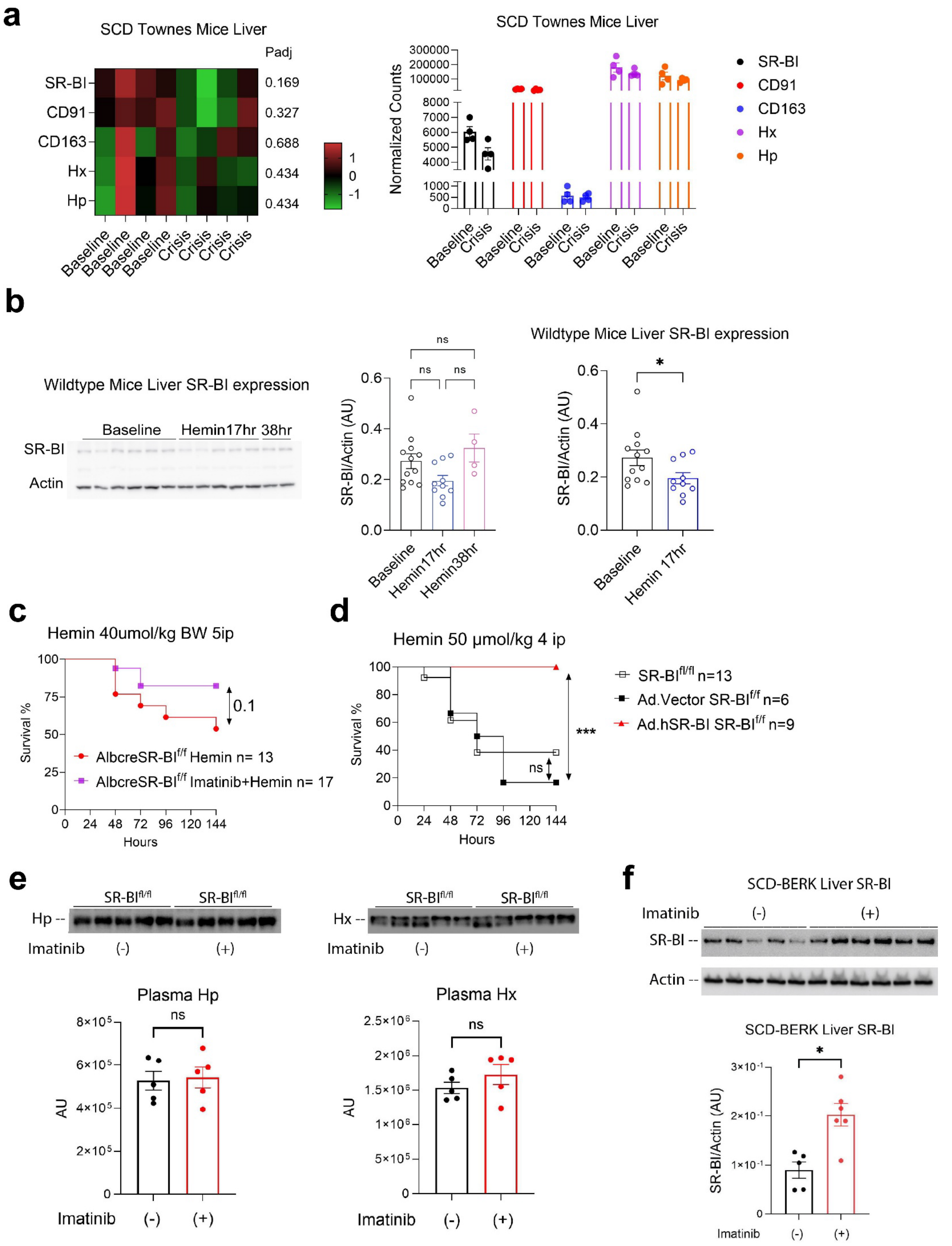
a
SCD Townes Mice Liver
Padj
SR-BI
CD91
CD163
Hx
Hp
0.169
0.327
0.688
0.434
0.434
1
0
-1
Baseline
Baseline
Baseline
Baseline
Crisis
Crisis
Crisis
Crisis
SCD Townes Mice Liver
Normalized Counts
300000
200000
100000
8000
7000
6000
5000
4000
1000
500
0
Baseline
Crisis
SR-BI
CD91
CD163
Hx
Hp
b
Wildtype Mice Liver SR-BI expression
Baseline
Hemin17hr
38hr
SR-BI
Actin
ns
ns
ns
SR-BI/Actin (AU)
0.6
0.4
0.2
0.0
Baseline
Hemin17hr
Hemin38hr
Wildtype Mice Liver SR-BI expression
*
Baseline
Hemin 17hr
c
Hemin 40umol/kg BW 5ip
Survival %
100
75
50
25
0
0.1
AlbcreSR-BIf/f Hemin n= 13
AlbcreSR-BIf/f Imatinib+Hemin n= 17
0 24 48 72 96 120 144
Hours
d
Hemin 50 μmol/kg 4 ip
Survival %
100
80
60
40
20
0
***
ns
SR-BIfl/fl n=13
Ad.Vector SR-BIf/f n=6
Ad.hSR-BI SR-BIf/f n=9
0 24 48 72 96 120 144
Hours
e
SR-BIfl/fl
SR-BIfl/fl
Hp --
Imatinib (-) (+)
SR-BIfl/fl
SR-BIfl/fl
Hx --
Imatinib (-) (+)
Plasma Hp
ns
8×10⁵
6×10⁵
4×10⁵
2×10⁵
0
AU
Imatinib (-) (+)
Plasma Hx
ns
2.5×10⁶
2×10⁶
1.5×10⁶
1×10⁶
5×10⁵
0
AU
Imatinib (-) (+)
f
SCD-BERK Liver SR-BI
Imatinib (-) (+)
SR-BI --
Actin --
SCD-BERK Liver SR-BI
*
3×10⁻¹
2×10⁻¹
1×10⁻¹
0
SR-BI/Actin (AU)
Imatinib (-) (+)

**Fig. S4.**

(**a**) Heatmap and normalized count of hemolysis scavenger gene expression at the liver of SCD Townes mice (Public Dataset). (**b**) SR-BI$^{fl/fl}$ (n = 4-12) mice were injected with 40 µmol/kg BW of hemin (i.p.). 30 µg of liver lysate were loaded and examined for SR-BI level via immunoblot. (**c**) Survival analysis of AlbcreSR-BI$^{fl/fl}$ mice treated with/without imatinib (50 mg/kg BW) for 3 days, followed by daily hemin injection. (**d**) Survival analysis of SR-BI$^{fl/fl}$ mice treat with no adenovirus, Ad.hSR-BI or Ad.vector for 3 days, followed by daily hemin injection. (**e**) Plasma Hp and Hx level in SR-BI$^{fl/fl}$ mice treated with/without imatinib for 3 days. 30 µg of plasma protein were loaded and examined for Hp and Hx via immunoblot. (**f**) SCD-BERK mice (n=5-6) were treated with or without imatinib (50 mg/kg BW) for 3 days. 30 µg of liver lysate were loaded and examined for SR-BI level via immunoblot. Survival curves were analyzed by the Log-Rank test. Significance was determined by two-way ANOVA for comparison across baseline, hemin 17 (early crisis) and 38hrs (late crisis) post injection and Mann-Whitney test for comparison between two groups. * $P < 0.05$; *** $P < 0.001$.
